# Pleiotropic win-win mutations can rapidly evolve in a nascent cooperative community despite unfavorable conditions

**DOI:** 10.1101/2020.07.21.214130

**Authors:** Samuel F. M. Hart, Chi-Chun Chen, Wenying Shou

## Abstract

Cooperation, paying a cost to benefit other individuals, is widespread. Cooperation can be promoted by pleiotropic “win-win” mutations which directly benefit self and partner. Previously, we showed that “partner-serving” should be defined as increased benefit supply rate per intake benefit (Hart & Pineda et al., 2019). Here, we report that “win-win” mutations can rapidly evolve even in nascent cooperation under conditions unfavorable for cooperation. Specifically, in a well-mixed environment we evolved engineered yeast cooperative communities where two strains exchanged costly metabolites lysine and hypoxanthine. Among cells that consumed lysine and released hypoxanthine, *ecm21* mutations repeatedly arose. *ecm21* is “self-serving”, improving self’s growth rate in limiting lysine. *ecm21* is also “partner-serving”, increasing hypoxanthine release rate per lysine consumption and the steady state growth rate of partner. *ecm21* also arose in monocultures evolving in lysine-limited chemostats. Thus, even without any pressure to maintain cooperation, pleiotropic win-win mutations may readily evolve.

## Introduction

Cooperation, paying a fitness cost to generate benefits available to others – is widespread and thought to drive major evolutionary transitions ^1,2^. For example in multi-cellular organisms, different cells must cooperate with each other and refrain from dividing in a cancerous fashion to ensure the propagation of the germline ^3^. Cooperation between species, or mutualistic cooperation, are also common ^4^. In extreme cases, mutualistic cooperation are obligatory, *i.e.* cooperating partners depend on each other for survival ^5,6^. For example, insects and endosymbiotic bacteria exchange costly essential metabolites ^6,7^.

Cooperation is vulnerable to “cheaters” who gain a fitness advantage over cooperators by consuming benefits without reciprocating fairly. Cancers are cheaters of multi-cellular organisms ^8^, and rhizobia variants can cheat on their legume hosts ^9^. How might cooperation survive cheaters?

Various mechanisms are known to protect cooperation against cheaters. In “partner choice”, an individual preferentially interacts with cooperating partners over spatially-equivalent cheating partners ^2,10–12^. For example, client fish observes cleaner fish cleaning other clients, and then chooses the cleaner fish that offers high-quality service (removing client parasites instead of client tissue) to interact with ^11^.

For organisms lacking partner choice mechanisms, a spatially-structured environment can promote the origin and maintenance of cooperation ^2,13–17^. This is because in a spatially-structured environment neighbors repeatedly interact, and thus cheaters will eventually suffer as their neighbors perish (“partner fidelity feedback”). In a well-mixed environment, since all individuals share equal access to the cooperative benefit regardless of their contributions, cheaters are favored over cooperators ^15^. An exception is that cooperators can stochastically purge cheaters if cooperators happen to be better adapted to an environmental stress than cheaters ^18–20^. Finally, pleiotropy, where a single mutation affects multiple phenotypes, can stabilize cooperation if reducing benefit supply to partner also elicits a crippling effect on self ^21–24^. For example, when the social amoeba *Dictyostelium discoideum* experience starvation, a fraction of the cells differentiate into a non-viable stalk in order to support the remaining cells to differentiation into viable spores. *dimA* mutants attempt to cheat by avoiding the stalk fate, but they also fail to form spores ^21^. As another example, during quorum sensing in *Pseudomonas aeruginosa*, cooperators pay a fitness cost to secret proteases that break down extracellular proteins into usable amino acids. *LasR* mutants that “cheat” by not secreting protease also fail to metabolize adenosine for themselves ^22^. In both cases, a gene, together with its associated network, links an individual’s cooperation capacity to the individual’s fitness, thus stabilizing cooperation. However, given the long evolutionary histories of these cooperative systems, it is unclear how easily such a genetic linkage can arise.

Here, we investigate whether cooperation-stabilizing “win-win” mutations could arise in a nascent cooperative community growing in an environment unfavorable for cooperation. The nascent cooperative community is termed CoSMO (<u>Co</u>operation that is <u>S</u>ynthetic and <u>M</u>utually <u>O</u>bligatory). CoSMO comprises two non-mating engineered *Saccharomyces cerevisiae* strains: *L*^−^*H*^+^ requires lysine (*L*) and pays a fitness cost to overproduce hypoxanthine (*H*, an adenine derivative) ^20,25^, while *H*^−^*L*^+^ requires hypoxanthine and pays a fitness cost to overproduce lysine ^26^ (Figure 1A). Overproduced metabolites are released into the environment by live cells ^25^, allowing the two strains to feed each other. CoSMO models the metabolic cooperation between certain gut microbial species ^27^ and between legumes and rhizobia ^28^, as well as other mutualisms ^29–34^.

**Figure 1.**
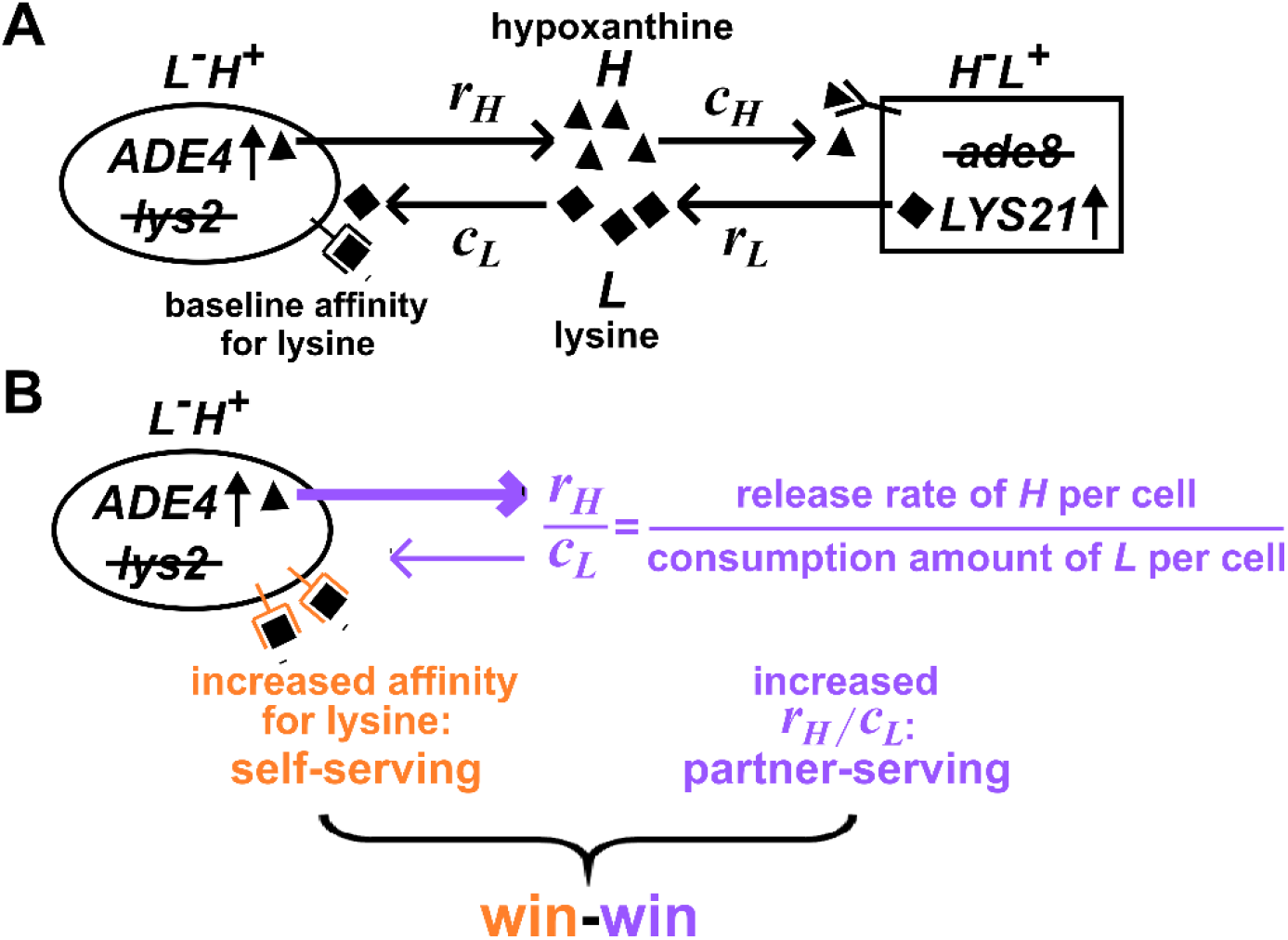
Win-win mutation in a nascent cooperative community. Figure 1 (**A**) CoSMO consists of two non-mating cross-feeding yeast strains, each engineered to overproduce a metabolite required by the partner strain. Metabolite overproduction is due to a mutation that renders the first enzyme of the biosynthetic pathway resistant to end-product inhibition ^47,48^. Hypoxanthine and lysine are released by live *L*^−^*H*^+^ and live *H*^−^*L*^+^ cells at a rate of *r*_*H*_ and *r*_*L*_ per cell, respectively ^25^, and are consumed by the partner at an amount of *c*_*H*_ and *c*_*L*_ per cell, respectively. The two strains can be distinguished by different fluorescent markers. (**B**) **Win-win mutation**. A win-win mutation is pleiotropic: it confers a self-serving phenotype (orange) and a partner-serving phenotype (lavender).

In our previous work, we allowed nine independent lines of CoSMO to evolve for over 100 generations in a well-mixed environment by performing periodic dilutions ^25,35^. Throughout evolution, the two cooperating strains coexisted due to their metabolic co-dependence ^35,36^. In a well-mixed environment, since partner-supplied benefits are uniformly distributed and equally available to all individuals, a self-serving mutation will be favored regardless of how it affects the partner. Indeed, all characterized mutants isolated from CoSMO displayed self-serving phenotypic changes ^20,25,26^, outcompeting their ancestor in community-like environments. Here, we report the identification of a pleiotropic win-win mutation which is both self-serving and partner-serving. This win-win mutation also arose in the absence of the cooperative partner. Thus, cooperation-promoting win-win mutations can arise in a community without any evolutionary history of cooperation and in environments unfavorable to cooperation.

## Results

### Criteria of a win-win mutation

A win-win mutation is defined as a single mutation (e.g. a point mutation; a translocation; a chromosome duplication) that directly promotes the fitness of self (“self-serving”) and the fitness of partner (“partner-serving”). To define “direct” here, we adapt the framework from Chapter 10 of (Peters et al., 2017): A mutation in genotype *A* exerts a direct fitness effect on genotype *B* if the mutation can alter the growth rate of *B* even if the biomass of *A* is fixed.

For *L*^−^*H*^+^, a self-serving mutation should improve the growth rate of self by, for example, increasing cell’s affinity for lysine (Figure 1B, orange). A self-serving mutation allows the mutant to outcompete a non-mutant. A partner-serving mutation should improve the growth rate of partner at a fixed self biomass. Since the partner requires hypoxanthine, a partner-serving mutation in *L*^−^*H*^+^ should increase the hypoxanthine supply rate per *L*^−^*H*^+^ biomass. Since the biomass of *L*^−^*H*^+^ is linked to lysine consumption, the partner-serving phenotype of *L*^−^*H*^+^ translates to hypoxanthine supply rate per lysine consumption, or equivalently, hypoxanthine release rate per cell (*r*_*H*_) normalized by the amount of lysine consumed to make a cell (*c*_*L*_) ^26^. We call this ratio “*H*-*L* exchange ratio” (Figure 1B, purple). Note that a partner-serving mutation will eventually feedback to promote self growth. Indeed, after an initial lag, the growth rate of partner, of self, and of the entire community reach the same steady state growth rate 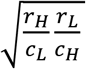, where *r*_*L*_ (lysine release rate per cell) and *c*_*H*_ (hypoxanthine consumption per cell) are phenotypes of *H*^−^*L*^+^ ^26^.

### Community and monoculture evolution share similar mutations

We randomly isolated evolved *L*^−^*H*^+^ colonies from CoSMO, and subjected them to whole-genome sequencing. Nearly every sequenced clone harbored one or more of the following mutations: *ecm21*, *rsp5*, and duplication of chromosome 14 (*DISOMY14*) (Table 1, top), consistent with our earlier studies ^20,25,26,37^. Mutations in *RSP5*, an essential gene, mostly involved point mutations (e.g. *rsp5*(*P772L*)), while mutations in *ECM21* mostly involved premature stop codons and frameshift mutations (Table 1, top; Table 1 Figure Supplement 1). Similar mutations also repeatedly arose when *L*^−^*H*^+^ evolved as a monoculture in lysine-limited chemostats (Table 1, bottom), suggesting that these mutations emerged independently of the partner.

**Table 1.**
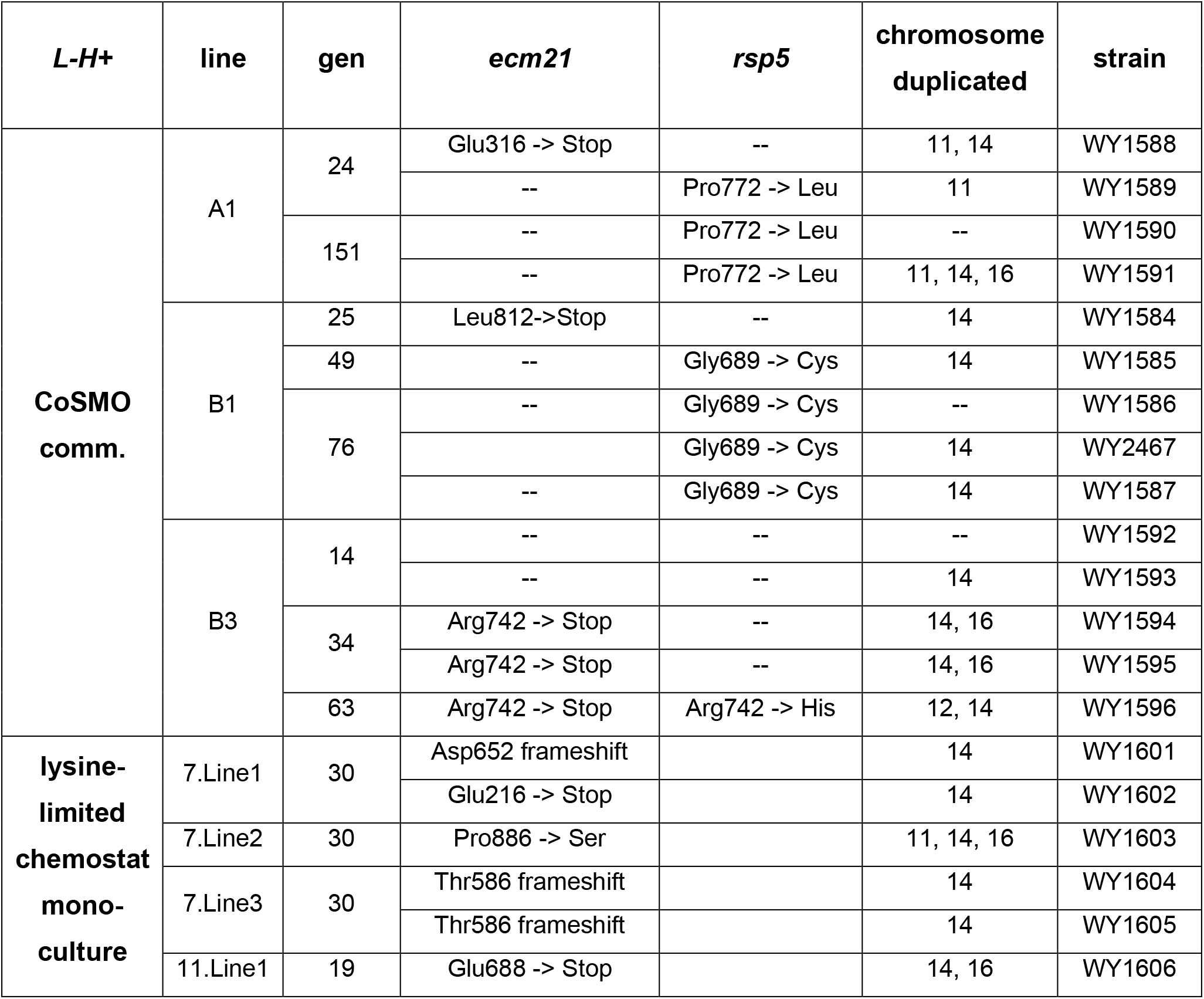
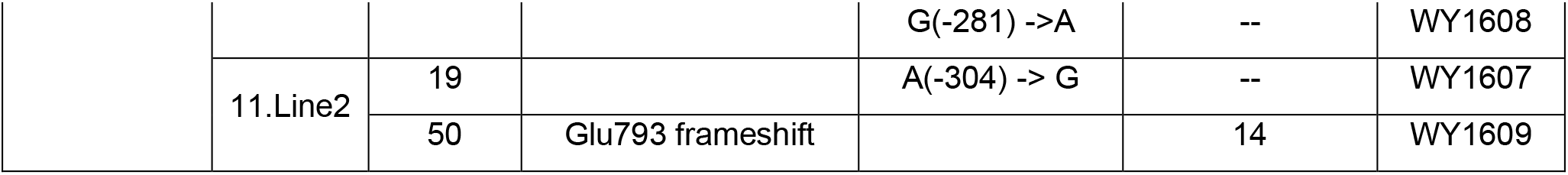
Mutations that repeatedly arose in independent lines. Table 1 Single-nucleotide polymorphisms (SNPs) and chromosomal duplications from Illumina re-sequencing of *L*^−^*H*^+^ from CoSMO communities (top) and lysine-limited chemostats (bottom). All clones except for two (WY1592 and WY1593 of line B3 at Generation 14) had either an *ecm21* or an *rsp5* mutation, often in conjunction with chromosome 14 duplication. Note that the RM11 strain background in this study differed from the S288C strain background used in our earlier study ^20^. This could explain, for example, why mutations in *DOA4* were repeatedly observed in the earlier study ^20^ but not here. For a schematic diagram of the locations of *ecm21* and *rsp5* mutations with respect to protein functional domains, see Table 1-Figure Supplement 1. For other mutations, see Table 1-Table Supplement 1.

### Self-serving mutations increase the abundance of lysine permease

Evolved *L*^−^*H*^+^ clones are known to display a self-serving phenotype: they could form microcolonies on low-lysine plates where the ancestor failed to grow ^20,26^. To quantify this self-serving phenotype, we used a fluorescence microscopy assay ^38^ to measure the growth rates of ancestral and evolved *L*^−^*H*^+^ in various concentrations of lysine. Under lysine limitation characteristic of the CoSMO environment (Figure 2A, “Comm. environ.”), evolved *L*^−^*H*^+^ clones containing an *ecm21* or *rsp5* mutation grew faster than a *DISOMY14* strain which, as we showed previously, grew faster than the ancestor ^39^. An engineered *ecm21Δ* or *rsp5*(*P772L*) mutation was sufficient to confer the self-serving phenotype (Figure 2A).

**Figure 2.**
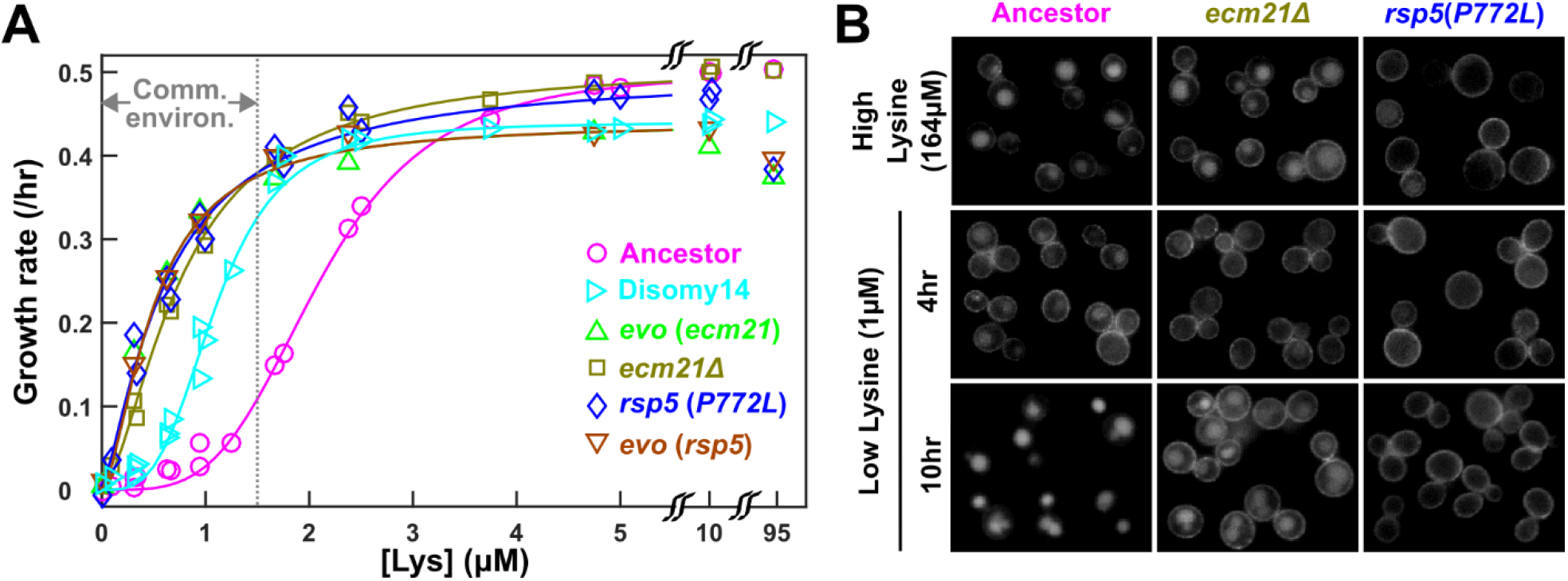
Self-serving mutations increase growth rates at low lysine via increasing membrane Lyp1. Figure 2 **(A) Recurrent mutations are self-serving**. We measured growth rates of mutant and ancestral strains in minimal SD medium with various lysine concentrations, using a calibrated fluorescence microscopy assay ^38^. Briefly, for each sample, total fluorescence intensity of image frames were tracked over time, and the maximal positive slope of ln(fluorescence intensity) against time was used as the growth rate. Evolved strains grew faster than the ancestor in community environment (the grey dotted line demarcating “Comm. environ.” corresponds to the lysine level supporting a growth rate of 0.1/hr as observed in ancestral CoSMO ^25^). Measurements performed on independent days (≥ 3 trials) were pooled and the average growth rate is plotted. Fit lines are based on Moser’s equation for the birth rate *b* as a function of metabolite concentration *L*: *b*(*L*) = *b*_max_ *L*^*n*^ / (*K*_*L*_^*n*^ + *L*^*n*^), where *b*_*max*_ is maximum birth rate in excess lysine, *K*_*L*_ is the lysine concentration at which half maximum birth rate is achieved, and *n* is the cooperitivity cooeficient describing the sigmoidal shape of the curve ^49^. Evolved strains are marked with “*evo*”; engineered or backcrossed mutants are marked with the genotype. Data for *DISOMY14* are reproduced from ^26^ as a comparison. Data can be found in “Figure 2 Source Data”. **(B) Self-serving mutations stabilize Lyp1 localization on cell membrane.** We fluorescently tagged Lyp1 with GFP in ancestor (WY1620), *ecm21Δ* (WY2355), and *rsp5*(*P772L*) (WY2356) to observe Lyp1 localization. We imaged each strain in a high lysine concentration (164 μM) as well as after 4 and 10 hours incubation in low lysine (1 μM). Note that low lysine was not consumed during incubation ^38^. During prolonged lysine limitation, Lyp1 was stabilized to cell membrane in both mutants compared to the ancestor.

The self-serving phenotype is due to an increased abundance of the high-affinity lysine permease Lyp1 on the cell membrane. We have previously shown that duplication of the *LYP1* gene, which resides on Chromosome 14, is necessary and sufficient for the self-serving phenotype of *DISOMY14* (Figure 2A) ^39^. Rsp5, an E3 ubiquitin ligase, is recruited by various “adaptor” proteins to ubiquitinate and target membrane transporters including Lyp1 for endocytosis and vacuolar degradation ^40^. In high lysine, Lyp1-GFP was localized to cell membrane and vacuole in ancestral and *ecm21* cells, but localized to the cell membrane in *rsp5* cells (Figure 2B, top row). Thus, Lyp1 localization was normal in *ecm21* but not in *rsp5*, consistent with the notion that at high lysine concentration, Lyp1 is targeted for ubiquitination by Rsp5 through the Art1 instead of the Ecm21 adaptor ^40^. When ancestral *L*^−^*H*^+^ was incubated in low lysine, Lyp1-GFP was initially localized on the cell membrane to facilitate lysine uptake, but later targeted to the vacuole for degradation and recycling ^41^ (Figure 2B bottom panels). However, in both *ecm21Δ* and *rsp5*(*P772L*) mutants, Lyp1-GFP was stabilized on cell membrane during prolonged lysine limitation (Figure 2B bottom panels), thus allowing mutants to grow faster than the ancestor during lysine limitation.

### ecm21 mutation is partner-serving

The partner-serving phenotype of *L*^−^*H*^+^ (i.e. hypoxanthine release rate per lysine consumption; exchange ratio *r*_*H*_/*c*_*L*_) can be measured in lysine-limited chemostats. In chemostats, fresh medium containing lysine was supplied at a fixed slow flow rate (mimicking the slow lysine supply by partner), and culture overflow exited the culture vessel at the same flow rate. After an initial lag, live and dead population densities reached a steady state (Figure 3- Figure supplement 1) and therefore, the net growth rate must be equal to the chemostat dilution rate *dil* (flow rate/culture volume). The released hypoxanthine also reached a steady state (Figure 3A). The *H*-*L* exchange ratio can be quantified as *dil* * *H*_*ss*_/*L*_0_ ^26^, where *dil* is the chemostat dilution rate, *H*_*ss*_ is the steady state hypoxantine concentration in the culture vessel, and *L*_*0*_ is the lysine concentration in the inflow medium (which was fixed across all experiments).

**Figure 3.**
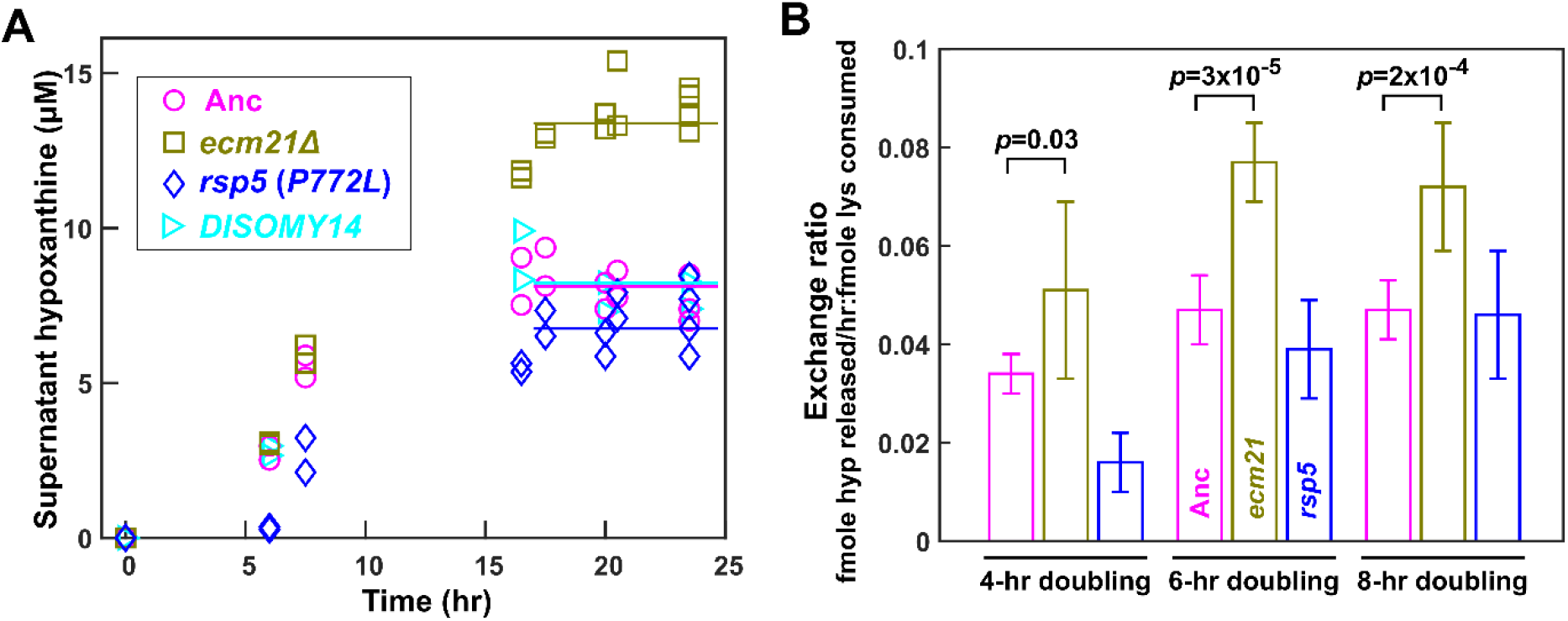
ecm21Δ releases more hypoxanthine per lysine consumption. Figure 3 **(A) Hypoxanthine accumulates to a higher level in *ecm21Δ* chemostats than in ancestor chemostats.** We cultured ancestor and mutant strains in lysine-limited chemostats (20 μM input lysine) at 6-hr doubling time (similar to CoSMO doubling time). Periodically, we quantified live and dead cell densities using flow cytometry (Figure 3 Figure Supplement 1), and hypoxanthine concentration in filtered supernatant using a yield-based bioassay ^25^. The steady state hypoxanthine concentration created by the ancestor (WY1335) was lower than e*cm21Δ* (WY2226), and slightly higher than *rsp5*(*P772L*) (WY2475). *DISOMY14* (WY2349) was indistinguishable from the ancestor, similar to our previous report ^26^. **(B) *ecm21Δ* has a higher hypoxanthine-lysine exchange ratio than the ancestor**. Cells were cultured in lysine-limited chemostats that spanned the range of CoSMO environments. In all tested doubling times, the exchange ratios of *ecm21Δ* were significantly higher than those of the ancestor. The exchange ratios of *rsp5*(*P772L*) are similar to or lower than those of the ancestor. Mean and two standard deviations from 4~5 experiments are plotted. *p*-values are from two-tailed t-test assuming either unequal variance (4-hr doubling) or equal variance (6-hr and 8-hr doublings; verified by F-test). Data and *p*-value calculations can be found in “Figure 3 Source Data”.

Compared to the ancestor, *ecm21*Δ but not *DISOMY14* ^26^ or *rsp5*(*P772L*) exhibited increased *H-L* exchange ratio. Specifically, at the same dilution rate (corresponding to 6-hr doubling), the steady state hypoxanthine concentration was the highest in *ecm21*Δ, and lower in the ancestor, *DISOMY14* ^26^, and *rsp5*(*P772L*) (Figure 3A). Although exchange ratio depends on doubling time, exchange ratios of *ecm21Δ* consistently outperformed those of the ancestor across doubling times typically found in CoSMO (Figure 3B). Thus, compared to the ancestor, *ecm21*Δ has a higher hypoxanthine release rate per lysine consumption.

To test whether *ecm21*Δ can promote partner growth rate, we quantified the steady state growth rate of the *H*^−^*L*^+^ partner when cocultured with either ancestor or *ecm21*Δ *L*^−^*H*^+^ in CoSMO communities. After the initial lag, CoSMO reached a steady state growth rate ^39^ (constant slopes in Figure 4A), which was also achieved by the two cooperating strains ^39^. Compared to the ancestor, *ecm21*Δ indeed sped up the steady state growth rate of CoSMO and of partner *H*^−^*L*^+^ (Figure 4B). Thus, *ecm21*Δ is partner-serving.

**Figure 4.**
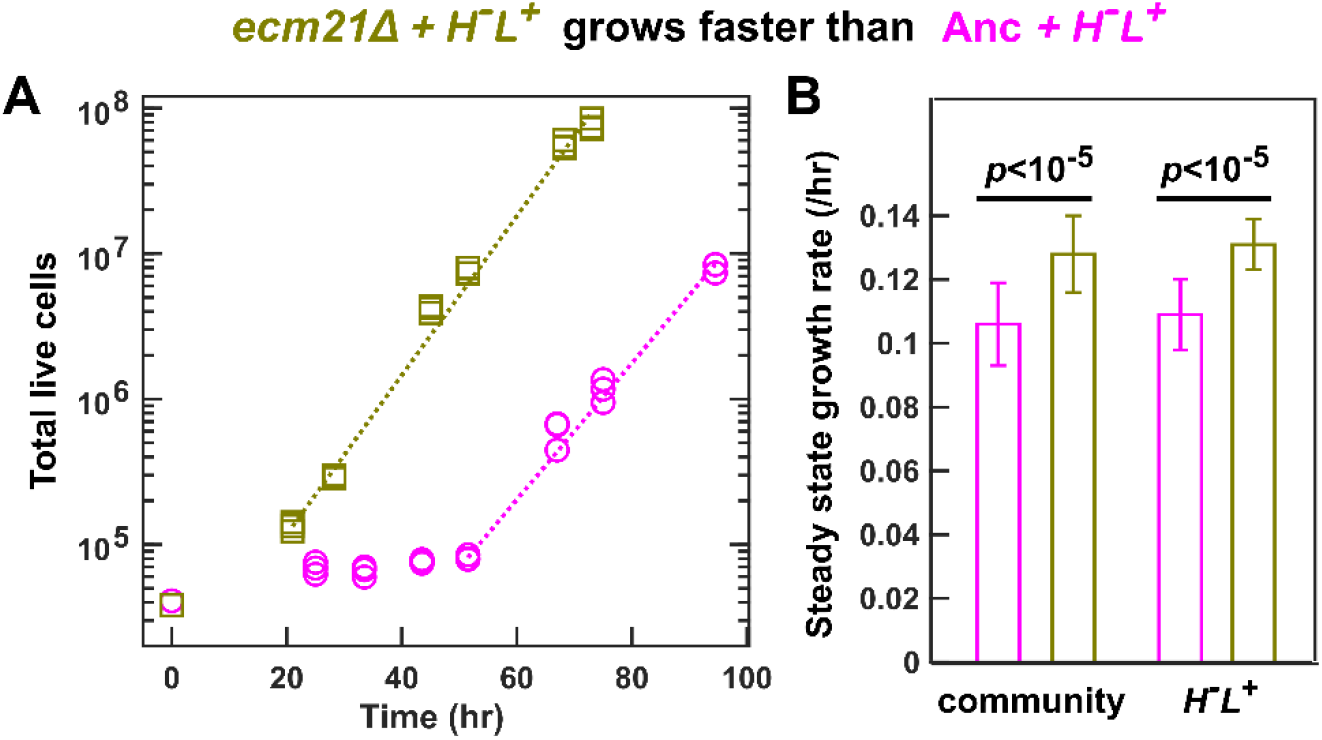
ecm21Δ increases the growth rate of CoSMO and of partner. To prevent rapid evolution, we grew CoSMO containing ancestral *H*^−^*L*^+^ and ancestral or *ecm21*Δ mutant *L*^−^*H*^+^ in a spatially-structured environment on agarose pads, and periodically measured the absolute abundance of the two strains using flow cytometry ^25^. (**A**) **Growth dynamics**. After an initial lag, CoSMO achieved a steady state growth rate (slope of dotted line). (**B**) ***ecm21Δ* increases the growth rate of CoSMO and of partner.** Steady state growth rates of the entire community (left) and of partner *H*^−^*L*^+^ (right) were measured (n≥6), and the average and two standard deviations are plotted. *p*-values are from two-tailed t-test with equal variance (verified by F-test). The full data set and outcomes of statistical tests can be found in Figure 4 Source Data.

The partner-serving phenotype of *ecm21*Δ can be explained by the increased hypoxanthine release rate per lysine consumption, rather than the evolution of any new metabolic interactions. Specifically, the growth rate of partner *H*^−^*L*^+^ (and of community) is approximately the geometric mean of the two strains’ exchange ratios, or 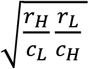 ^25,39^. Here, the ancestral partner’s exchange ratio 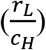 is fixed, while the exchange ratio of *L*^−^*H*^+^ 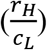 is ~1.6-fold increased in *ecm21*Δ compared to the ancestor (at doubling times of 6~8 hrs; Figure 3B). Thus, *ecm21*Δ is predicted to increase partner growth rate by 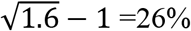 (95% confidence interval: 12%~38%; Figure 3 Source Data). In experiments, *ecm21*Δ increased partner growth rate by ~21% (Figure 4B; Figure 4 Source Data).

In conclusion, when *L*^−^*H*^+^ evolved in nascent mutualistic communities and in chemostat monocultures in a well-mixed environment, win-win *ecm21* mutations repeatedly arose (Table 1). Thus, pleiotropic win-win mutations can emerge in the absence of any prior history of cooperation, and in environments unfavorable for cooperation.

## Discussions

Here, we demonstrate that pleiotropic win-win mutations can rapidly arise in conditions unfavorable for cooperation – such as in a well-mixed environment or in the absence of the cooperative partner. As expected, all evolved *L*^−^*H*^+^ clones displayed self-serving phenotypes, achieving a higher growth rate than the ancestor in low lysine via stabilizing the lysine permease Lyp1 on cell membrane (Figure 2). Surprisingly, *ecm21* mutants also displayed partner-serving phenotypes, promoting the steady state growth rate of partner *H*^−^*L*^+^ and of community (Figure 4) via increasing the hypoxanthine release rate per lysine consumption (Figure 3).

The partner-serving phenotype of *L*^−^*H*^+^ emerged as a side-effect of adaptation to lysine limitation instead of adaptation to a cooperative partner. We reached this conclusion because *ecm21* mutations were also observed in *L*^−^*H*^+^ evolving as monocultures in lysine-limited chemostats (Table 1). Being self-serving does not automatically lead to a partner-serving phenotype. For example in the *DISOMY14* mutant, duplication of the lysine permease *LYP1* improved mutant’s affinity for lysine (Figure 2) without improving hypoxanthine release rate per lysine consumption (Figure 3A) or partner’s growth rate ^26^.

How might *ecm21* achieve higher hypoxanthine release rate per lysine consumption? One possibility is that purine overproduction is increased in *ecm21* mutants, leading to a steeper concentration gradient across the cell membrane. A different, and not mutually exclusive, possibility is that in *ecm21* mutants, purine permeases are stabilized much like the lysine permease, which in turn leads to an increased membrane permeability. Future work will reveal mechanisms of win-win mutations, as well as how common win-win mutations might be.

Here, we have defined a win-win mutation as a pleiotropic mutation that directly promotes the fitness of self and the fitness of partner. A broader definition of partner-serving is also possible if we include mutations that *indirectly* promote partner’s fitness through promoting self fitness. Consider a mutant with improved affinity for lysine but no alterations in the metabolite exchange ratio (e.g. *DISOMY14*). By growing better in low lysine, this mutant will improve its own survival which in turn helps the whole community (and thus the partner) to survive the initial stage of low cell density. Indeed, all evolved *L*^−^*H*^+^ clones tested so far improved community (and partner) viability in the sense that all mutants reduced the minimal total cell density required for the community to grow to saturation (Shou et al., 2007; Waite and Shou, 2012). Unlike *ecm21*, some of these mutations (e.g. *DISOMY14*) are not directly partner-serving, and would not improve partner’s steady state growth rate ^26^.

How likely might cooperation be stabilized by pleiotropy? Given that gene networks display “small world” connectivity ^42^ and that a protein generally interacts with many other proteins, mutations in one gene will likely affect multiple cellular processes. Indeed, pleiotropy has been found to stabilize several natural cooperative systems ^21–24^. In all these known examples, cooperation is intra-population, and has a long evolutionary history of cooperation. Our work demonstrates that pleiotropy can give rise to win-win mutations that promote nascent, mutualistic cooperation. The emergence of win-win mutations may not require elaborate evolutionary history, or selective pressure to maintain cooperation (e.g. in a well-mixed environment and even in the absence of the cooperative partner). In the absence of a cooperative partner, an abiotic environment could still sometimes mimic the community environment due to, for example, the presence of detritus. Adaptation to this abiotic environment may sometimes select for win-win mutations that will later benefit an incoming partner. Along with these earlier works, our study highlights the possibility of pleiotropy influencing the origin and maintenance of mutualistic cooperation.

## Methods

### Strains

Our nomenclature of yeast genes, proteins, and mutations follows literature convention. For example, the wild type *ECM21* gene encodes the Ecm21 protein; *ecm21* represents a reduction-of loss-of-function mutation. In our evolution experiments, we used *S. cerevisiae* strains of the RM11-1a background instead of the original S288C background ^35^, because the latter had a significantly higher frequency of generating petites with defective mitochondria and compromised DNA-repair capability ^43^. Both *L*^−^*H*^+^ (WY1335) and *H*^−^*L*^+^ (WY1340) were of *MATa* and harbored the *ste3Δ* mutation (Table S1). Absence of the a-factor receptor Ste3 would prevent mating between the two strains even after a rare event that switches a cell from *MATa* to *MATalpha*. We introduced desired genetic modifications into the ancestral RM11 background via transformation ^44,45^. Strains were stored at −80°C in 15% glycerol. All evolved or engineered strains used in this paper are summarized in Table 1 (evolved clones) and Supplementary File 1 (ancestral and engineered strains).

Growth medium and strain culturing have been previously discussed ^26^.

### Experimental evolution

CoSMO evolution has been described in detail in ^26^. Briefly, exponentially-growing *L*^−^*H*^+^ (WY1335) and *H*^−^*L*^+^ (WY1340) were washed free of supplements, counted using a Coulter counter, and mixed at 1000:1 (Line A), 1:1 (Line B), or 1:1000 (Line C) at a total density of 5×10^5^/ml. Three 3ml community replicates (replicates 1, 2, and 3) per initial ratio were initiated, thus constituting nine independent lines. Since the evolutionary outcomes of the nine lines were similar, they could be treated as a single group. Communities were grown at 30°C in glass tubes on a rotator to ensure well-mixing. Community turbidity was tracked by measuring the optical density (OD_600_) in a spectrophotometer once to twice every day. In this study, 1 OD was found to be 2~4×10^7^cells/ml. We diluted communities periodically to maintain OD at below 0.5 to avoid additional selections due to limitations of nutrients other than adenine or lysine. The fold-dilution was controlled to within 10~20 folds to minimize introducing severe population bottlenecks. Coculture generation was calculated from accumulative population density by multiplying OD with total fold-dilutions. Sample were periodically frozen down at −80°C. To isolate clones, a sample of frozen community was plated on rich medium YPD and clones from the two strains were distinguished by their fluorescence colors or drug resistance markers.

For chemostat evolution of *L*^−^*H*^+^, device fabrication and setup are described in detail in ^37^. Briefly, the device allowed the evolution of six independent cultures, each at an independent doubling time. To inoculate each chemostat vessel, ancestral *L*^−^*H*^+^ (WY1335) was grown to exponential phase in SD supplemented with 164 μM lysine. The cultures were washed with SD and diluted to OD600 of 0.1 (~7×10^6^/ml) in SD. 20 ml of diluted culture was added to each vessel through the sampling needle, followed by 5 ml SD to rinse the needle of excess cells. Of six total chemostat vessels, each containing ~43mL running volume, three were set to operate at a target doubling time of 7 hours (flow rate ~4.25 mL/hr), and three were set to an 11 hour target doubling time (flow rate ~2.72 mL/hr). With 21 μM lysine in the reservoir, the target steady state cell density was 7×10^6^/ml. In reality, live cell densities varied between 4×10^6^/ml and 1.2×10^7^/ml. Samples were periodically taken through a sterile syringe needle. The nutrient reservoir was refilled when necessary by injecting media through a sterile 0.2 micron filter through a 60-ml syringe. We did not use any sterile filtered air, and were able to run the experiment without contamination for 500 hours. Some reservoirs (and thus vessels) became contaminated after 500 hours.

Whole-genome sequencing of evolved clones and data analysis were described in detail in ^26^.

### Quantification methods

Microscopy quantification of *L*^−^*H*^+^ growth rates at various lysine concentrations was described in ^25,38^. Briefly, cells were diluted into flat-bottom microtiter plates to low densities to minimize metabolite depletion during measurements. Microtiter plates were imaged periodically (every 0.5~2 hrs) under a 10x objective in a temperature-controlled Nikon Eclipse TE-2000U inverted fluorescence microscope. Time-lapse images were analyzed using an ImageJ plugin Bioact ^38^. We normalized total fluorescence intensity against that at time zero, calculated the slope of ln(normalized total fluorescence intensity) over three to four consecutive time points, and chose the maximal value as the growth rate corresponding to the input lysine concentration. For validation of this method, see ^38^.

Short-term chemostat culturing of *L*^−^*H*^+^ for measuring exchange ratio was described in ^25,38^. Briefly, because *L*^−^*H*^+^ rapidly evolved in lysine-limited chemostat, we took special care to ensure the rapid attainment of steady state so that an experiment is kept within 24 hrs. We set the pump flow rate to achieve the desired doubling time *T* (19ml culture volume*ln(2)/*T*). Periodically, we sampled chemostats to measure live and dead cell densities, and the concentration of released hypoxanthine.

Cell density measurement via flow cytometry was described in ^25^. Briefly, we mixed into each sample a fixed volume of fluorescent bead stock whose density was determined using a hemocytometer or Coulter counter. From the ratio between fluorescent cells or non-fluorescent cells to beads, we can calculate live cell density and dead cell density, respectively.

Chemical concentration measurement was performed via a yield-based bioassay ^25^. Briefly, the hypoxanthine concentration in an unknown sample was inferred from a standard curve where the final turbidities of an *ade-* tester strain increased linearly with increasing concentrations of input hypoxanthine.

Quantification of CoSMO growth rate was described in ^25^. Briefly, we used the “spot” setting where a 15 μl drop of CoSMO community (1:1 strain ratio; ~4×10^4^ total cells/patch) was placed in a 4-mm inoculum radius in the center of a 1/6 Petri-dish agarose sector. During periodic sampling, we cut out the agarose patch containing cells, submerged it in water, vortexed for a few seconds, and discarded agarose. We then subjected the cell suspension to flow cytometry.

### Imaging of GFP localization

Cells were grown to exponential phase in SD plus 164μM lysine. A sample was washed with and resuspended in SD. Cells were diluted into wells of a Nunc 96-well Optical Bottom Plate (Fisher Scientific, 165305) containing 300μl SD supplemented with 164μM or 1μM lysine. Images were acquired under a 40X oil immersion objective in a Nikon Eclipse TE2000-U inverted fluorescence microscope equipped with a temperature-controlled chamber set at 300C. GFP was imaged using an ET-EYFP filter cube (Exciter: ET500/20×, Emitter: ET535/30m, Dichroic: T515LP). Identical exposure times (500 msec) were used for both evolved and ancestral cells.

### Introducing mutations into the essential gene RSP5

Since *RSP5* is an essential gene, the method of deleting the gene with a drug-resistance marker and then replacing the marker with a mutant gene cannot be applied. We therefore modified a two-step strategy ^46^ to introduce a point mutation found in an evolved clone into the ancestral *L*^−^*H*^+^ strain. First, a loxP-kanMX-loxP drug resistance cassette was introduced into ~300 bp after the stop codon of the mutant *rsp5* to avoid accidentally disrupting the remaining function in *rsp5*. Second, a region spanning from ~250 bp upstream of the point mutation [C(2315)→T] to immediately after the loxP-kanMX-loxP drug resistance cassette was PCR-amplified. The PCR fragment was transformed into a wild-type strain lacking *kanMX*. G418-resistant colonies were selected and PCR verified for correct integration (11 out of 11 correct). The homologous region during transformation is large, and thus recombination can occur in such a way that the transformant got the *KanMX* marker but not the mutation. We therefore Sanger-sequenced the region, found that 1 out of 11 had the correct mutation, and proceeded with that strain.

---

**Figure 3 Figure Supplement 1.**
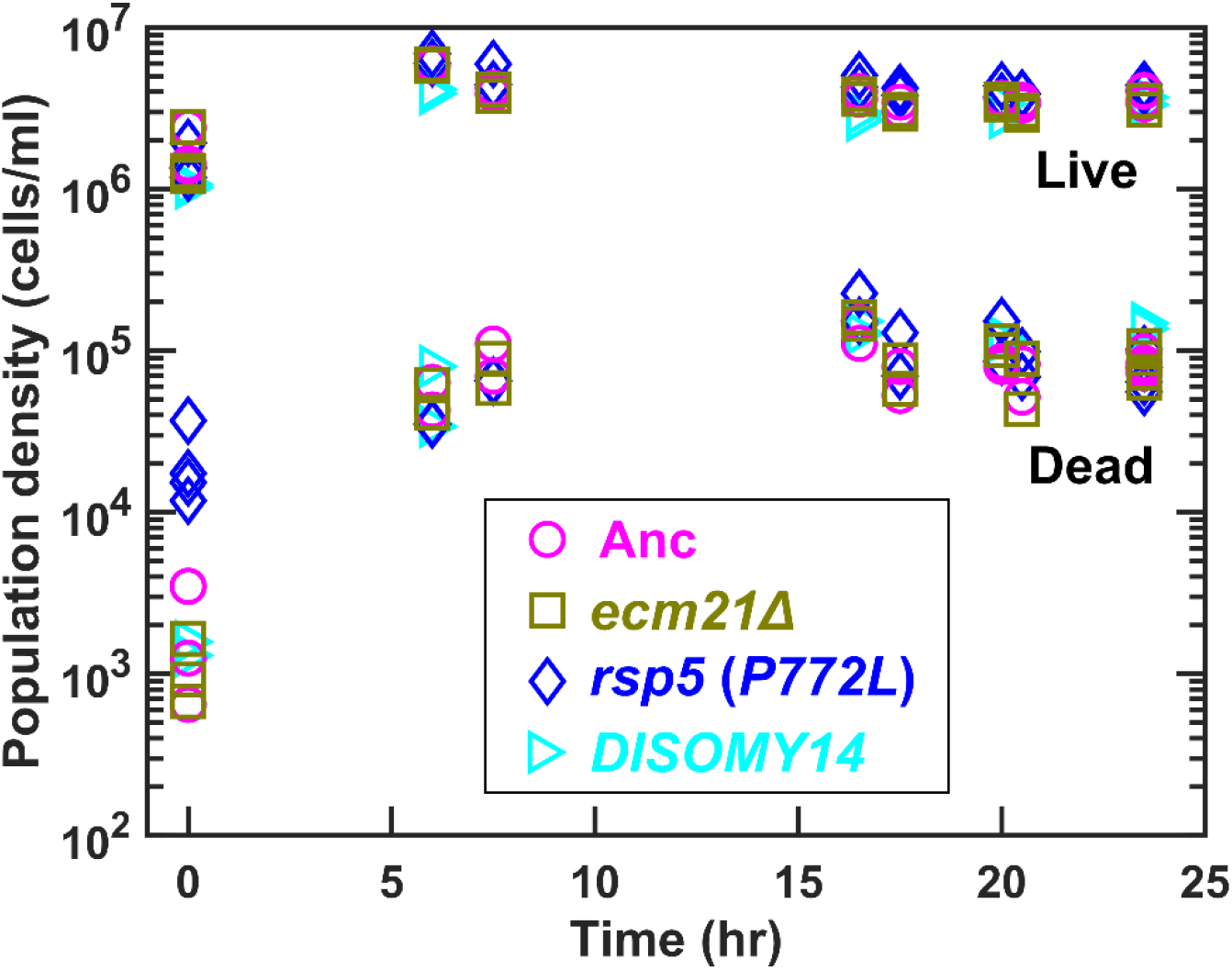
We cultured ancestor and mutant strains in lysine-limited chemostats (20 μM input lysine) at 6-hr doubling time (similar to CoSMO doubling time). Periodically, we measured live and dead cell densities using flow cytometry ^25^. After a lag, live and dead cell densities reached a steady state. Data can be found in “Figure 3 Source Data”.

**Table 1-Figure Supplement 1.**
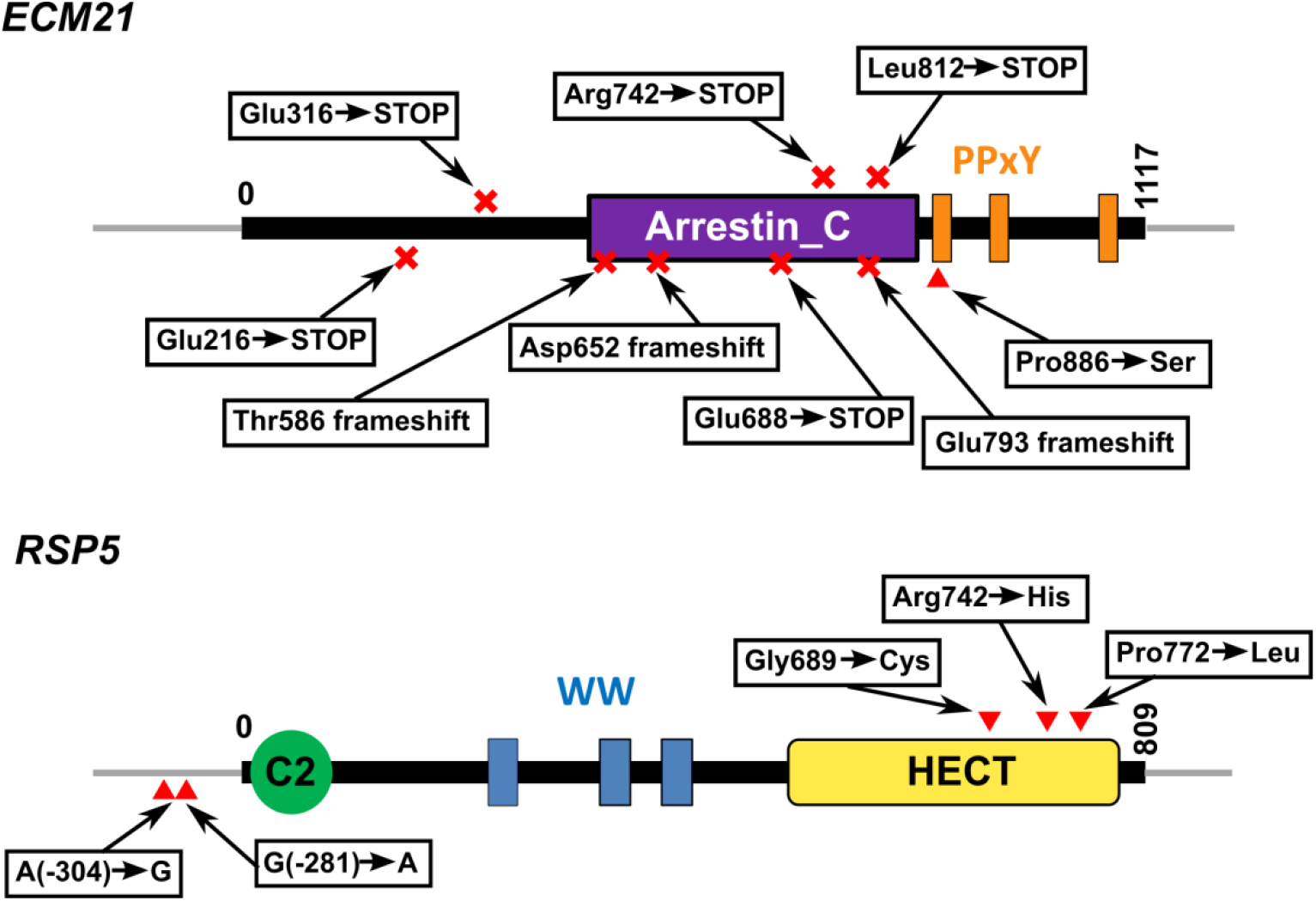
Functional domains and positions of mutations in Ecm21 and Rsp5 proteins. Mutations and their locations are marked with respect to the functional domains of the proteins. Numbers indicate amino acid positions, except in non-coding regions. Doman structures are obtained from the “protein” tab of SGD (https://www.yeastgenome.org/locus/S000000927/protein; https://www.yeastgenome.org/locus/S000000197/protein). HECT domain is found in ubiquitin-protein ligases. WW domain can bind proteins with particular proline-motifs such as the PPxY motif. Arrestin C-terminal-like domain is involved in signaling and endocytosis of receptors. For *ECM21*, mutating the three poly-proline-tyrosine (PY) motifs after amino acid 884 inhibited the stress-induced endocytosis of the manganese transporter Smf1 ^50^. Most *ecm21* mutations we recovered introduced premature stop codons before the PY motifs. In *RSP5*, the region including and upstream of - 470 is required for *RSP5* function ^51^. Mutations from coculture and monoculture isolates are marked above and below the gene, respectively.

**Table 1-Table Supplement 1. Summary of mutations**

**Supplementary file 1. List of strains**

## Acknowledgement

We thank Aric Capel for the chemostat evolution experiment data, and Jose Pineda for an earlier collaboration that eventually led to this discovery. We also thank members of the Shou lab (Li Xie, David Skelding, Alex Yuan, Sonal) for discussions.

